# SIRT4 enhances the cytotoxicity of NK cells toward hepatic stellate cells and reverses liver fibrosis via AMPKα/P-p53/NKG2DL pathway

**DOI:** 10.1101/2024.09.30.615768

**Authors:** Huan Chen, Binlin Da, Zihao Cai, Rui Fang, Xiaolin Xie, Han Zhang, Si Zhao, Ming Zhang, Lei Wang, Bing Xu, Yuzheng Zhuge, Feng Zhang

## Abstract

Natural killer (NK) cells exhibit antifibrotic properties in liver fibrosis (LF) by suppressing activated hepatic stellate cells (HSCs). SIRT4, a mitochondrial regulatory protein, plays a crucial role as a link between energy metabolism and cell viability. However, the role of SIRT4 in the cytotoxicity of NK cells toward HSCs remains unexplored.

In this study, we found that SIRT4 was markedly downregulated in both mouse models and patients with LF. The loss of SIRT4 reduced the cytotoxicity of NK cells against activated HSCs, while its overexpression enhanced this cytotoxicity. Mechanistically, SIRT4 activates AMPKα to promote p53 phosphorylation and facilitates its nuclear translocation, which induces the transcription of ULBP1 and ULBP2, members of the NK group 2D Legend (NKG2DL) family of molecules. Finally, overexpression of SIRT4 activated mouse hepatic NK cells and reversed LF by constructing adeno-associated viruses (AAV) that specifically target HSCs.

Thus, SIRT4 is essential for the cytotoxicity of NK cells toward HSCs, and AAV8-pGAFP-SIRT4 may serve as a therapeutic approach for managing LF.

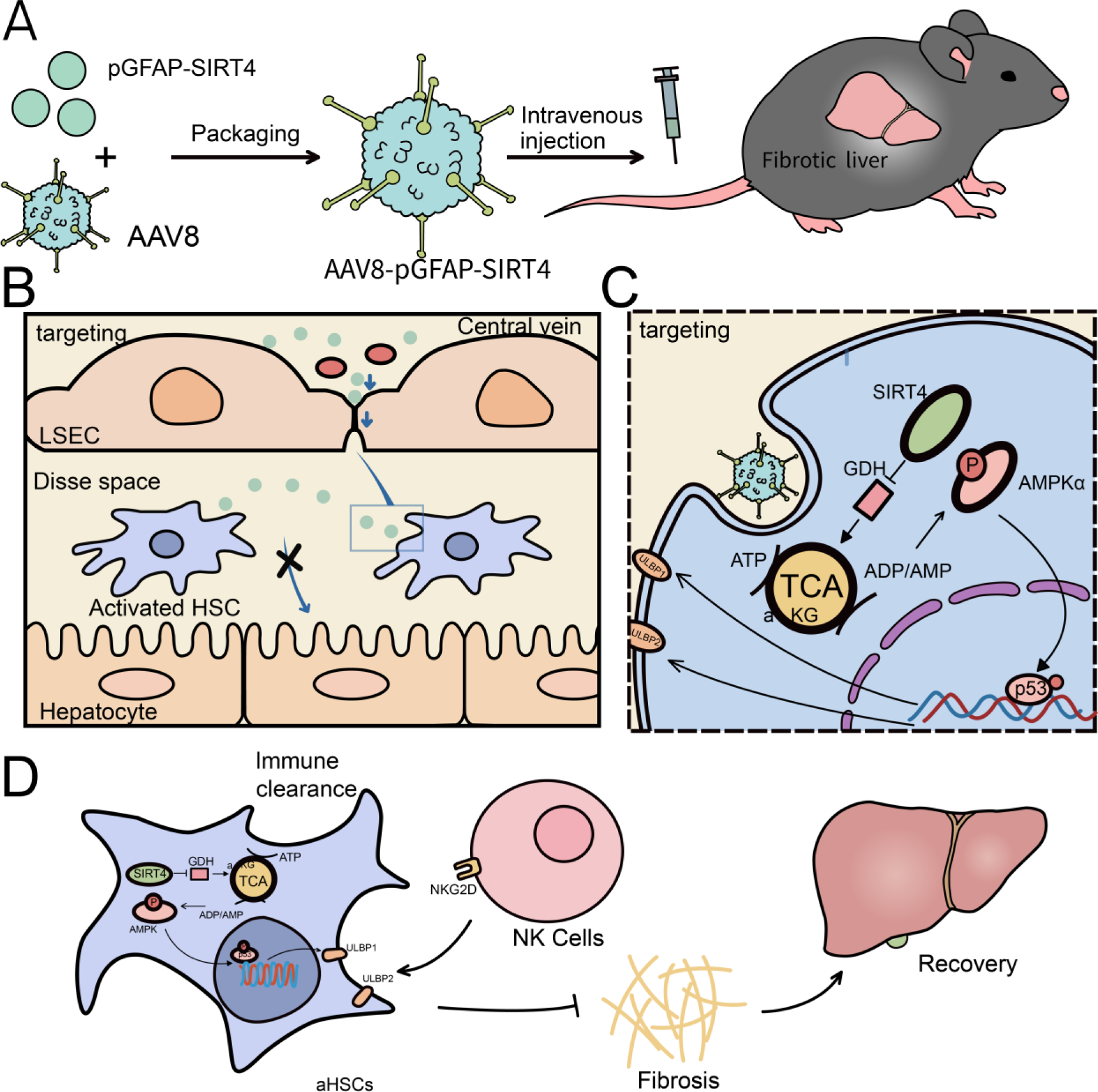

## Introduction

Chronic liver disorders (CLDs) continue to be serious illnesses that affect lives and are costly to society. The primary factor contributing to morbidity and death in patients with CLDs is liver fibrosis(LF)[1]. The exact processes of LF underlying some etiologies are yet unknown. The activation of hepatic stellate cells (HSCs) is essential for many different biological functions in LF. For instance, the loss of hepatic parenchymal structure and the upregulation and deposition of extracellular matrix (ECM)-related proteins produced by activated HSCs are characteristics of LF [2]. Since no proven therapy for LF has been found yet, strategies to reverse it have become crucial priorities.

Natural killer (NK) cells are an integral part of innate immunity in the liver and are vital in eliminating pathogens such as bacteria, viruses, and cancer cells[3]. Through techniques like genetic engineering, adoptive cell transfer of allogeneic NK cells, NK cell-targeted chemotherapy, and others, manipulation of NK cell activation has emerged as a promising cancer immunotherapy[4]. According to several earlier assessments, NK cells have antifibrotic properties that are enhanced by their cytotoxicity against activated HSCs and the release of other cytokines[5]. Consequently, NK cells activation is thought to be a potential treatment approach for LF[6]. The relationship between NK cells and HSCs is affected by a variety of stimulatory and inhibitory receptors located on NK cells, along with the respective ligands present in HSCs. Among the activating receptors identified on NK cells, NK group 2D (NKG2D) and NKp46 have been the focus of extensive research in the context of liver fibrosis. Previous studies indicate that the expression of NKG2D ligands (NKG2DL), such as major histocompatibility complex (MHC) class I polypeptide-related sequence A (MICA) and UL16 binding proteins (ULBPs), are elevated in senescent HSCs, rendering them more vulnerable to NK cell-mediated destruction via granule exocytosis. These ligands interact with NKG2D receptors on NK cells, triggering their activation, which results in the release of various cytotoxic agents and ultimately leads to the apoptosis of HSCs[7–8].

SIRT4, a mitochondria-localized sirtuin, controls the breakdown of many nutrients, including fat and glutamine, as well as the oxidation of fatty acids[9–11]. However, the role of SIRT4 in liver fibrosis was previously unknown. We for the first time found that SIRT4 expression was reduced in activated HSCs in vitro and vivo, and overexpression of SIRT4 significantly inhibited the proliferation of HSCs [12]. The results of recently published studies also showed that SIRT4 expression was decreased in the liver of patients with LF, and ex vivo experiments confirmed that SIRT4 was able to exert an antifibrotic effect through down-regulation of the Smads signaling pathway[13–14]. These results demonstrate that SIRT4 plays an important role in the reversal of LF, but the available data mainly reveal the effect of SIRT4 n HSCs, while the role and mechanism of removing activated HSCs are very poorly explored.In this study, we found that SIRT4 in HSCs plays a pivotal role in regulating the function of NK cells. Our findings reveal that SIRT4 drives the activation of, which modifies p53. This modification allows the p53 protein to enter the nucleus and function as a transcription factor, thereby increasing the transcription of ULBP1 and ULBP2. This process facilitates the recognition and elimination of HSCs by NK cells.

Adeno-associated virus (AAV) gene therapy has emerged as an effective and long-lasting gene therapy approach in recent years[15]. AAV vectors are often used for the lifelong treatment of genetic defects, with their therapeutic efficacy validated in multiple clinical trials [16]. AAVs with organ specificity can avoid side effects such as toxicity and immunogenicity associated with gene editing [17]. AAV8 is a serotype that targets the liver. In the present study, we constructed the AAV8 that specifically targets HSCs and assessed the efficacy of AAV8 as a vector for delivering pGFAP-SIRT4 into target HSCs against LF[18].

In summary, we observed that SIRT4 regulates the function of NK cells through the AMPKα/P-p53/NKG2DL signaling axis. The AAV8-GFAP-SIRT4 effectively overexpresses SIRT4 in HSCs and reverses LF, indicating its potential as a novel therapeutic approach for LF.

## Results

### The expression of SIRT4 is decreased in activated HSCs (aHSCs) and fibrotic livers

To investigate the potential role of SIRT4 in liver fibrosis, we first extracted primary mouse HSCs. Freshly isolated HSCs were considered quiescent HSCs (qHSCs), and when cultured for 7 days, qHSCs become aHSCs. With the onset of HSC activation, the protein expression of SIRT4 was greatly decreased (Figure 1A). Meanwhile, in mouse models of liver fibrosis induced by CCl4 or DDC, the expression of SIRT4 was also markedly reduced (Figure 1B, 1C). Consistently, the hepatic expression of SIRT4 was decreased in patients with liver fibrosis (Figure 1D). Taken together, these results suggest that SIRT4 may play a pivotal role in HSC activation and liver fibrosis.

**Fig. 1.**
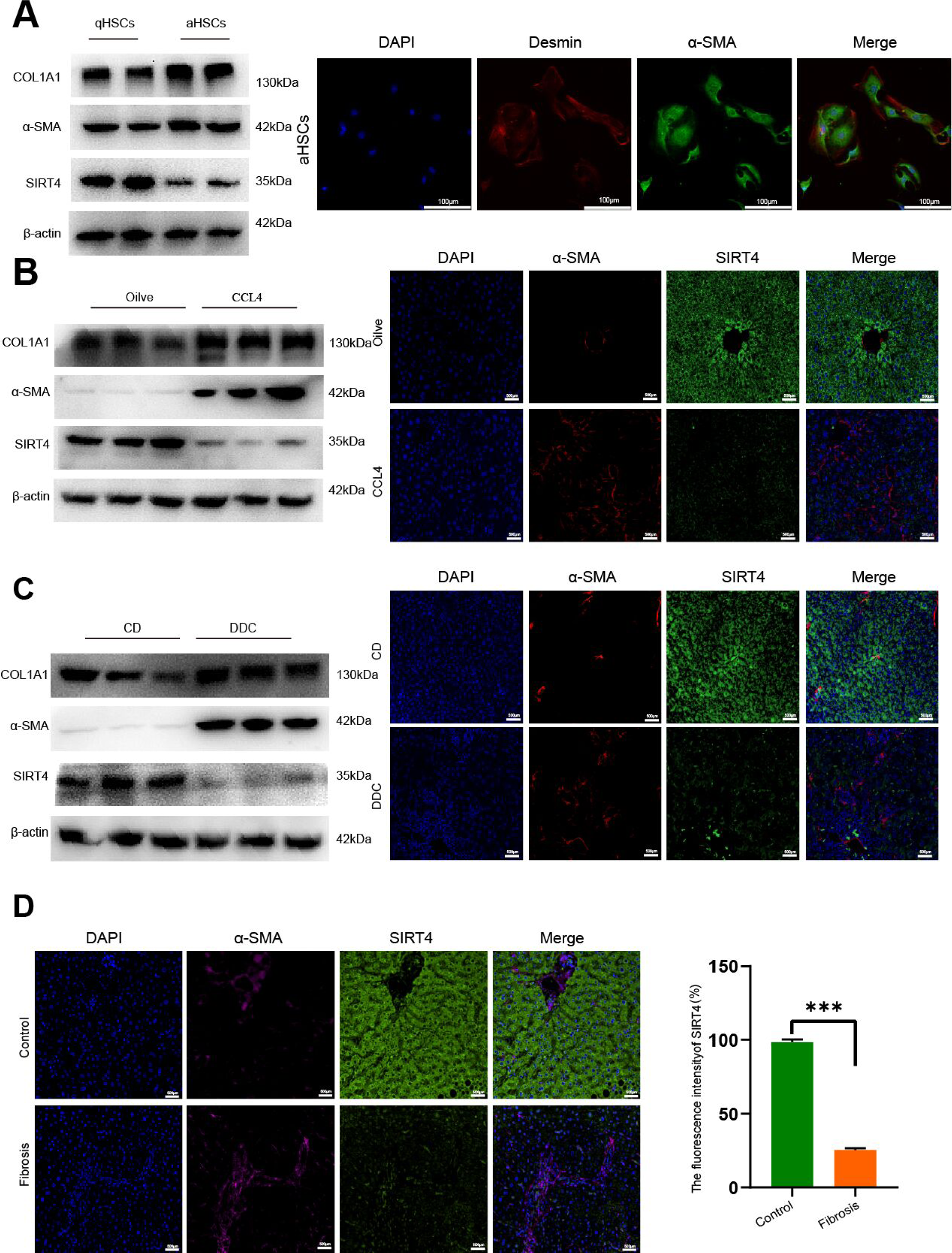
The expression of SIRT4 was decreased in aHSCs and fibrotic livers of mice and humans. (A) Primary HSCs were isolated from WT mice and cultured for 1 day (qHSCs) or 7 days (aHSCs). SIRT4 expression was decreased during primary HSC activation. (B) SIRT4 expression was decreased in the fibrotic liver of CCl4-treated mice. SIRT4 expression were detected by western blot analysis, immunofluorescence staining (C) SIRT4 expression was decreased in fibrotic liver of mice induced by DDC. (D) Liver sections were collected from normal individuals or patients with liver fibrosis and stained with Sirt4 and α-SMA. Right panel: quantification of Sirt4 fluorescent signal. Data are mean ± SEM. \**P* < 0.05, \*\**P* < 0.01,***P<0.001. Abbreviation: DAPI.

### Overexpression of SIRT4 in HSCs increased the cytotoxicity of NK cells

In our previous study, we described the protective role of SIRT4 in the pathogenesis of liver fibrosis. SIRT4 controls GDH enzyme activity and expression, regulating glutamine metabolism to inhibit HSCs proliferation. Choi et al [19] found that the uptake of glutamate was increased in activated HSCs and that mGluR5 activation enhanced the cytotoxicity of NK cells. Thus, we aim to investigate whether SIRT4 plays a role in regulating NK cell function during liver fibrosis. We overexpressed SIRT4 in LX-2 cells by transfecting plasmids (Figure 2A). To observe the elimination activity of NK cells against HSCs, NK92 cells (a human NK cell line) were used to co-culture with LX-2 cells with or without SIRT4 overexpression at different T: E ratios. Flow cytometry assay revealed a significant increase in the proportion of NK cells CD107a-positive cell population after overexpression of SIRT4 (Figure 2B). Meanwhile, we applied LDH cytotoxicity assay and found that the killing of LX-2 cells by NK92 cells was significantly increased after overexpression of SIRT4 (Figure 2C). Finally, we applied Calcein AM and PI probes to detect LX-2 cells after co-culture suggesting that overexpression of SIRT4 significantly increased the proportion of apoptotic cells in LX-2 cells (Figure 2D). In culture LX-2 cells, SIRT4 overexpression greatly increased the mRNA expression of ULBP1 and ULBP2, but neither MICA nor MICB was significantly altered (Figure 2E).

**Fig. 2.**
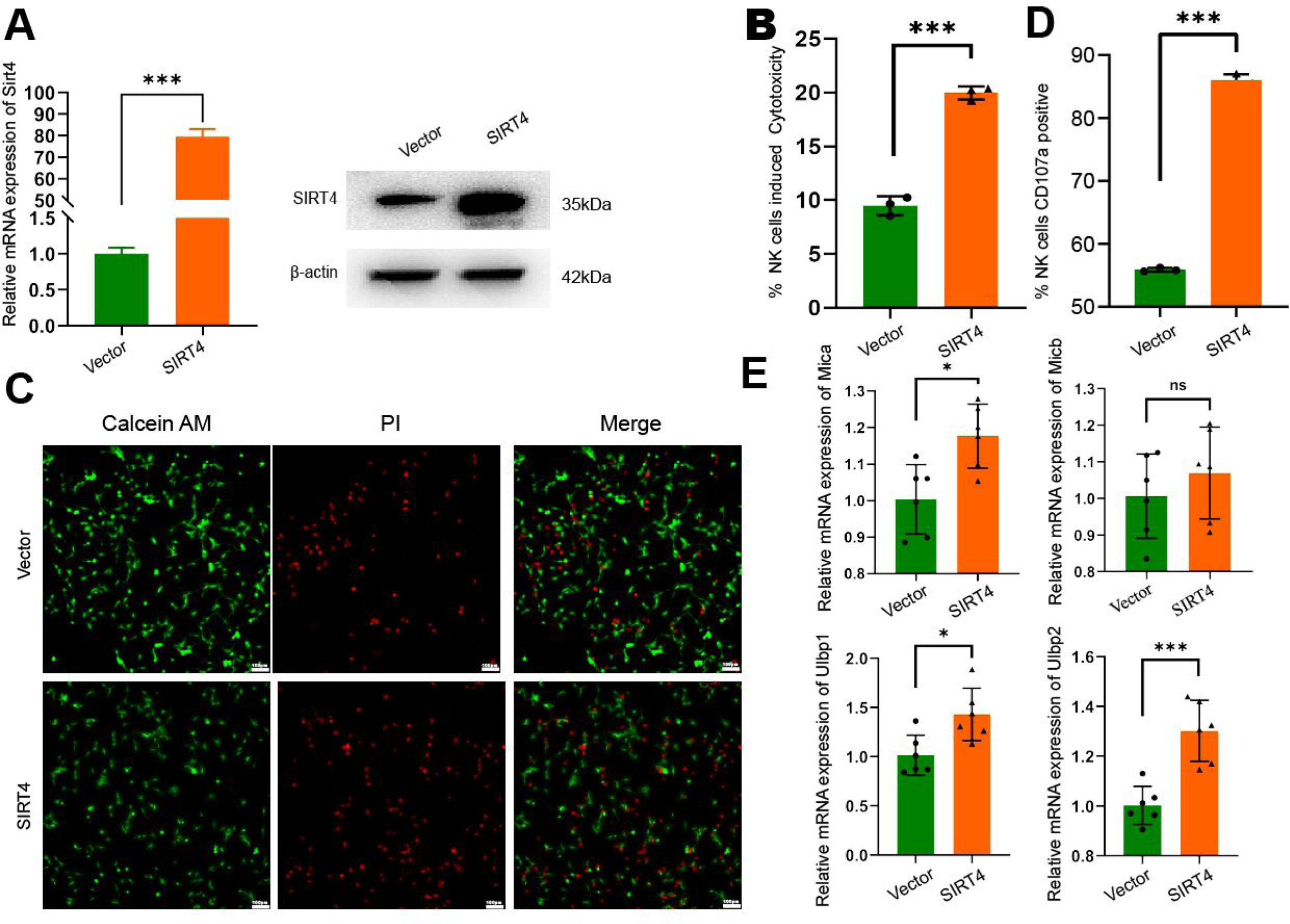
Overexpression of SIRT4 in HSCs increased the cytotoxicity of NK cells. (A)Overexpression of the SIRT4 gene in LX-2 cells with plasmids. QPCR and western blot analysis detection of overexpression efficiency (B)NK92 cells and LX-2 cells were co-cultured at a ratio of 10:1,for 6h, LDH cytotoxicity assay was used to detech NK cytotoxic effects (C)NK92 cells and LX-2 cells were co-cultured at a ratio of 10:1 for 6h, immunofluorescence analysis was used to detech apoptosis of LX-2 cells. (D) NK92 cells and LX-2 cells were co-cultured in a 10:1 ratio for 6 hours and showed upregulation of CD107a expression in NK92 cells as detected by flow cytometry (E) Overexpression of Sirt4, ULBP1,ULBP2 expression of NKG2DL protein family in LX-2 cells was upregulated by QPCR and western blot analysis \**P* < 0.05, \*\**P* < 0.01,***P<0.001.

### Knockdown of SIRT4 in LX-2 reduced the cytotoxicity of NK cells

To demonstrate the role of SIRT4 more fully, we constructed the LX-2 knockdown SIRT4 cell line by infection with lentivirus (Figure 3A). Further, co-culture experiments demonstrated a decrease in the proportion of NK92-activated cell numbers after the knockdown of SIRT4(Figure 3B). The LDH cytotoxicity assay responds to the attenuated killing toxicity of NK92 on LX-2(Figure 3C). Finally, we tested and found that the proportion of apoptosis in LX-2 cells was down-regulated in co-culture experiments after the knockdown of SIRT4 by Calcein AM and PI probes(Figure 3D). QPCR assays revealed downregulation of ULBP1 and ULBP2, but not MICA and MICB, expression after knockdown of SIRT4(Figure 3 E).

**Fig. 3.**
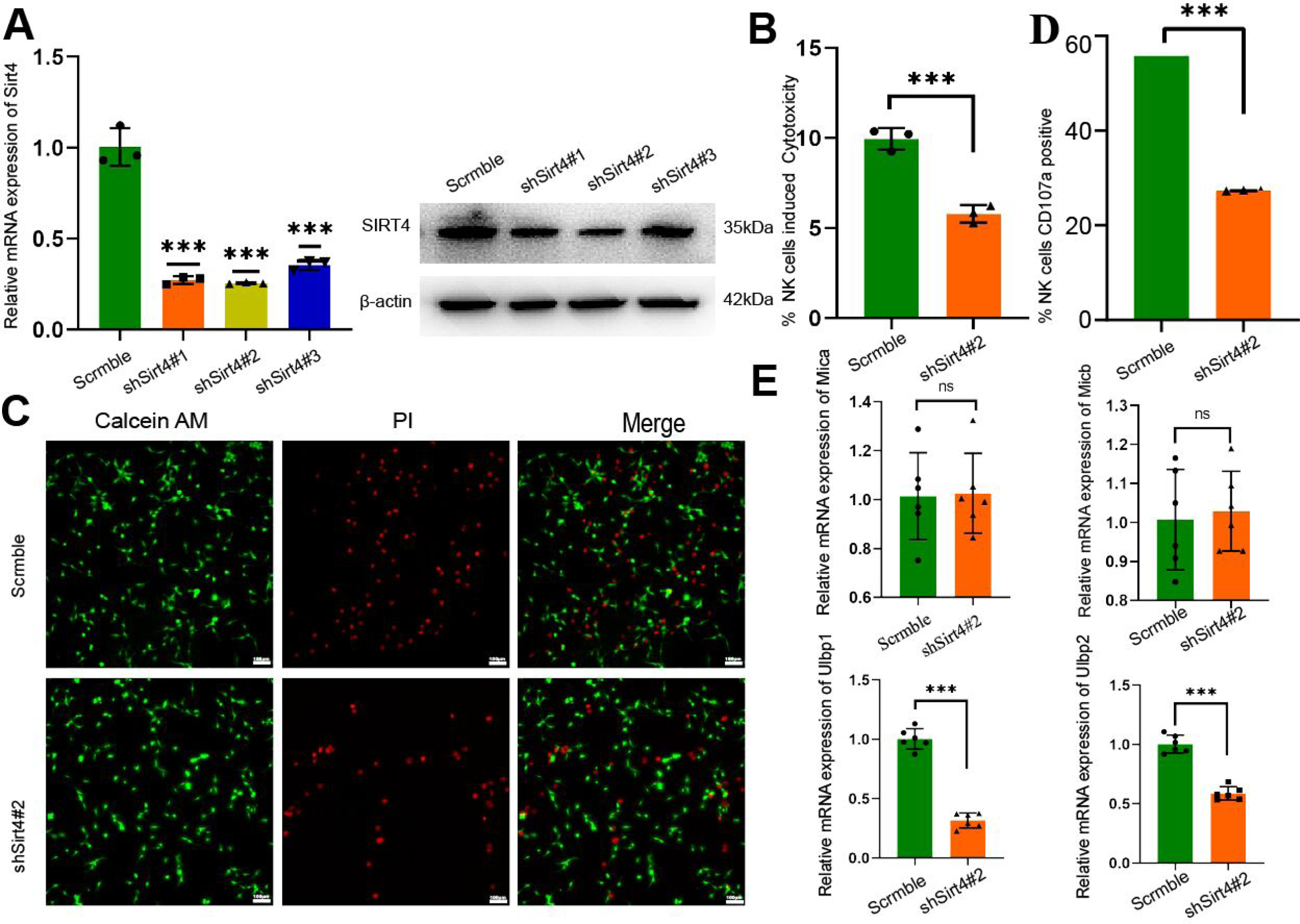
Knockdown of SIRT4 in HSCs decreased the the cytotoxicity of NK cells. (A) Knockdown of the Sirt4 gene in LX-2 cells with shSIRT4. QPCR and western blot analysis detection of knockdown efficiency (B) NK92 cells and LX-2 cells were co-cultured at a ratio of 10:1for 6h, LDH cytotoxicity assay was used to detech NK cytotoxic effects (C)NK92 cells and LX-2 cells were co-cultured at a ratio of 10:1 for 6h, immunofluorescence analysis was used to detech apoptosis of LX2. (D)NK92 cells and LX-2 cells were co-cultured in a 10:1 ratio for 6 hours and showed upregulation of CD107a expression in NK92 cells as detected by flow cytometry (E)Knockdown of SIRT4, MICA,MICB,ULBP1,ULBP2 expression of NKG2DL protein family in LX-2 cells was upregulated by QPCR. \**P* < 0.05, \*\**P* < 0.01,***P<0.001.

### SIRT4 mediates p53 phosphorylation by regulating AMPKα activation

To further clarify how SIRT4 induced NK cytotoxic effects, we performed RNA sequencing and analyzed the gene expression profile in the LX-2 cells with or without SIRT4 Knockdown. KEGG pathway enrichment analysis showed that SIRT4 was involved in regulating the AMPK signaling pathway and p53 signaling pathway (Figure 4A, 4B). We next experimentally explored whether SIRT4 was involved in regulating the p53 signaling pathway. We found that overexpression of SIRT4 reduced the p53 phosphorylation but had no effect on the total p53(Figure 4C, 4D).. Moreover, contradictory trends were found when SIRT4 was knockdown (Figure 4E, 4F). The previous study has found that SIRT4 can activate AMPKα by regulating glutamine metabolism and ultimately inhibit hepatocarcinogenesis [20]. We aimed to determine whether SIRT4 played the same role in HSCs. We found that the ectopic expression of SIRT4 significantly activated AMPKα (Figure 5A). Consistently, depletion of SIRT4 significantly reduced the phosphorylation of AMPKα (Figure 5B). Also after the knockdown of SIRT4 followed by the application of BETES (10μM), an agonist of glutamine metabolism, we found that the expression of phosphorylated AMPKα was up-regulated, suggesting that SIRT4 regulated the activation of AMPKα by inhibiting glutamine metabolism (Figure 5C). We knockdown SIRT4 in LX2 cells before using metformin, an activator of AMPKα, and found that metformin reversed the decrease in phosphorylation of p53 brought about by the knockdown of SIRT4(Figure 5D, 5E).

**Fig. 4.**
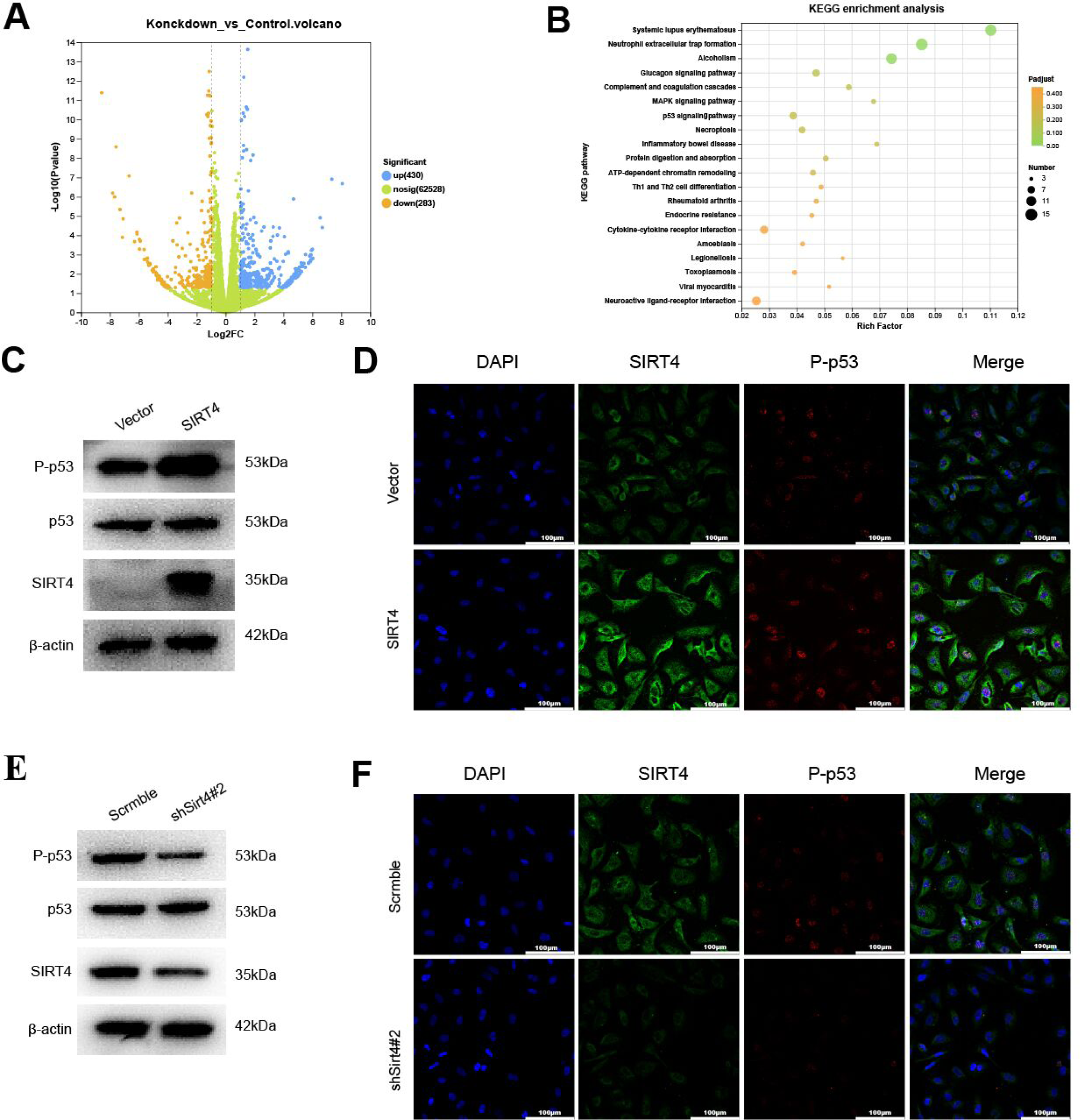
SIRT4 mediates p53 phosphorylation in HSCs. (A) (B)The RNA-seq and KEGG analysis of the differentially expressed genes in LX2 cell with or without SIRT4 Knockdown. (C)(D) Western blotting analysis and immunofluorescence was used to detect the phosphorylation level of p53 protein and p53 protein after overexpression of SIRT4 in LX2 cells. (E)(F) Western blotting analysis and immunofluorescence was used to detect the phosphorylation level of p53 protein and p53 protein after knockdown SIRT4 in LX2

**Fig 5.**
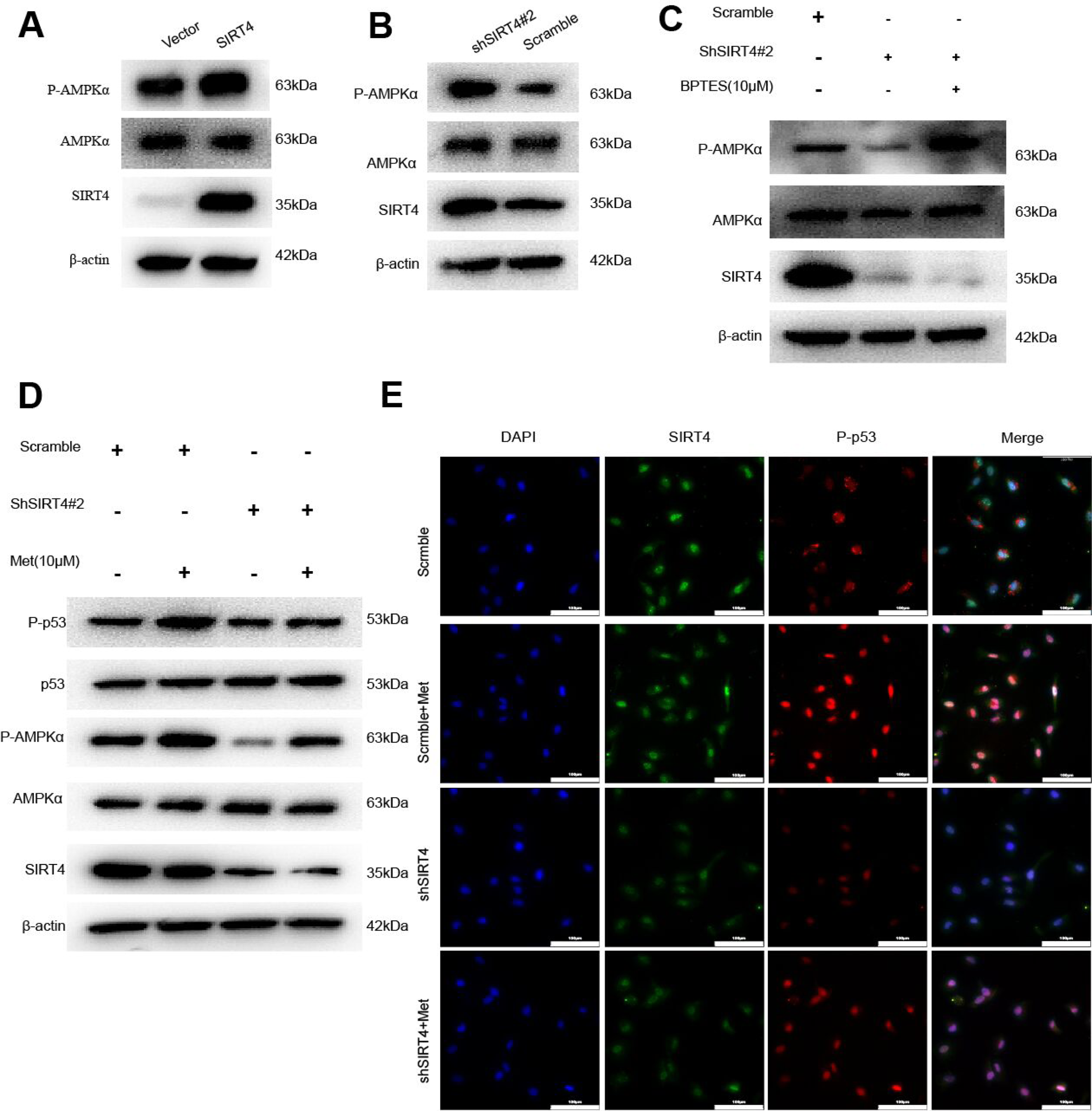
SIRT4 regulates AMPKα and p53 phosphorylation. (A) (B)Western blotting analysis wasused to detect the phosphorylation level of AMPKα protein and AMPKα protein after knockdown or overexpression SIRT4 in LX2 cells. (C) LX2 cells with or without BPTES(10μM) after knockdown SIRT4, detect the phosphorylation level of AMPKα protein and AMPK protein by using western blotting analysis. (D) (E)LX2 cells with or without metformin after knockdown SIRT4, detect the phosphorylation level of AMPKα protein, AMPK protein, the phosphorylation level of p53 protein and p53 protein by using western blotting analysis and immunofluorescence analysis.

### SIRT4 mediates ULBP1/ULBP2 transcriptional activation via the phosphorylation of p53 protein

To demonstrate that p53 is an important target of Sirt4 to enhance NK cytotoxicity, SIRT4 was knocked down in LX-2 followed by the application of Inauhzin, an agonist of p53. The result showed that the expression of ULBP1/ULBP2 was inhibited by SIRT4 knockdown, which was back-complemented by the ehancement of p53 (Figure 6A,6B).To investigate how p53 regulates ULBP1/ULBP2 expression, we first predicted whether the p53 protein has a binding site to the promoter region of ULBP1/ULBP2 on the transcription factor prediction website Jasper. The results suggest that p53 has a binding site with the promoter region of ULBP1/ULBP2, which suggests that p53 may act as a promoter of ULBP1/ULBP2 to enhance its transcription(Figure 6C). By CUT&RUN-qPCR, it was suggested that p53 was enriched for promoter DNA fragments of ULBP1/ULBP2, and the enrichment was more pronounced after overexpression of SIRT4(Figure 6D). Dual luciferase assay also showed that p53 promotes reporter gene expression through the ULBP1/ULBP2 promoter(Figure 6E). Meanwhile, the results of co-culturing the above-treated cells with NK92 cells suggested that the killing effect of NK92 on LX2 was attenuated after knockdown of SIRT4, and this effect was weakened by re-administration of Inauhzin(Figure 6F,6G,6H).

**Fig 6.**
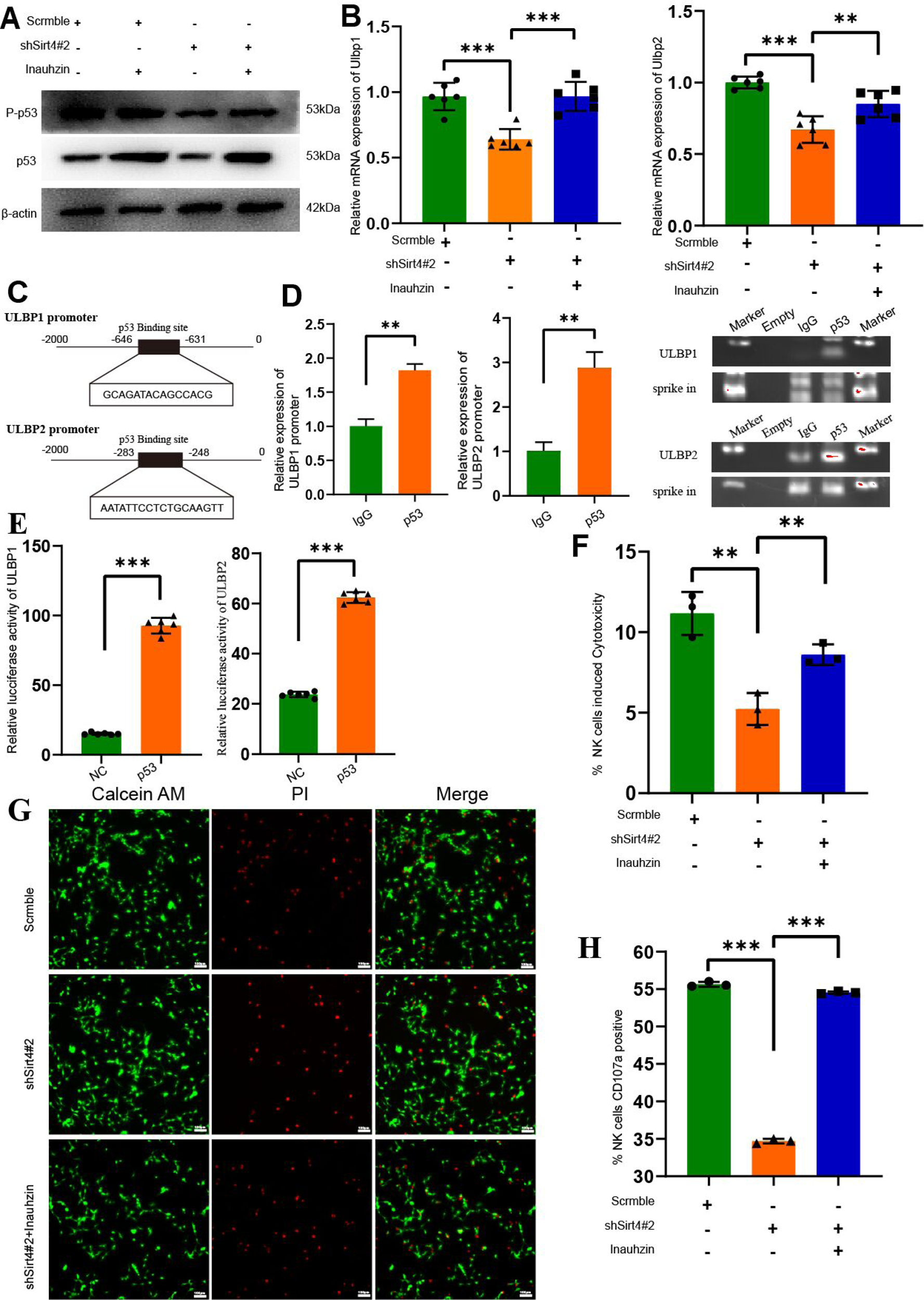
P-p53 plays an key role in the regulation of NK cytotoxicity by SIRT4. Agonist application of p53 after knockdown SIRT4 (A)Detecting the level of p53, P-p53(ser15) and SIRT4 by using Western blotting analysis. (B)QPCR was used to detect the mRNA expression of ULBP1/ULBP2.(C)Predicted promoter binding region of p53 to ULBP1/ULBP2 by Jasper (D)CUT&RUN was used to detect P-p53 binding to predicted sites (E)The luciferase assay was used to assess P53 binding to the ULBP1/ULBP2 promoter to drive reporter gene transcription (F)NK92 cells and LX-2 cells were co-cultured at a ratio of 5:1,10:1,20:1 for 6h, LDH cytotoxicity assay was used to detech NK cytotoxic effects (G)NK92 cells and LX-2 cells were co-cultured at a ratio of 10:1 for 6h, immunofluorescence analysis was used to detech apoptosis of LX-2 cells (H)NK92 cells and LX-2 cells were co-cultured in a 10:1 ratio for 6 hours and showed upregulation of CD107a expression in NK92 cells as detected by flow cytometry. \**P* < 0.05, \*\**P* < 0.01,***P<0.001.

### The HSC-specific overexpression of SIRT4 attenuates LF

Because HSC activation acts as a crucial contributor to LF, we hypothesized that therapies targeting SIRT4 in HSCs may protect against liver fibrosis. Similar to previous studies, the liver-specific gene delivery of AAV8 containing the promoter of glial fibrillary acidic protein (GFAP) was employed for targeted SIRT4 overexpression in HSCs in mice to validate this hypothesis. As shown in Figure 7A, Figure 7B, and Figure 7C, the successful delivery of exogenous SIRT4 into HSCs was confirmed by staining of Desmin-positive cells. At the same time, we examined the activation indexes of mouse liver NK cells, suggesting that the percentage of NK cell activation was elevated after overexpression of SIRT4(Figure 7D). In conclusion, HSCs-specific overexpression of SIRT4 enhances the toxic killing of HSCs by NK cells and reverses LF. As expected, SIRT4 was specifically overexpressed in HSCs, which significantly alleviated liver injury and fibrosis, as indicated by improvements in liver H&E, Masson, Sirius Red, and α-SMA staining and the levels of fibrogenic and HSCs activation-related markers (Figure 7E; Figure 7F). In addition, LF in mice undergoes another exacerbation when scavengers of NK cells are administered after delivery of SIRT4.

**Fig. 7.**
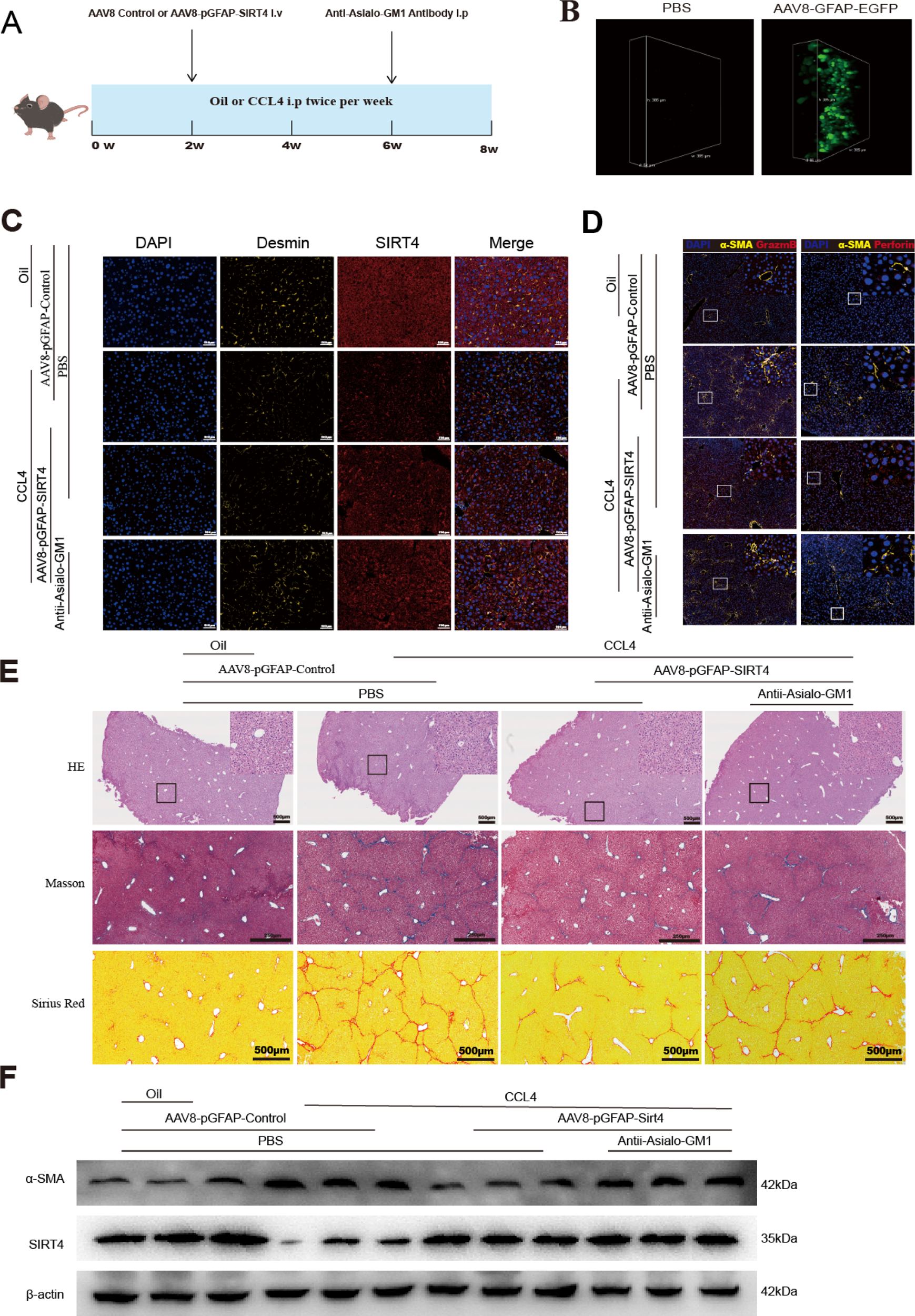
Overexpression of SIRT4 via HSCs-specific AAV8 inverse liver fibrosis. The HSC-specific overexpression of Sirt4 was induced by the injection of AAV8-pGFAP-Sirt4 and Antii-Asialo-GM1 treatment into mice exposed to CCl4. (A) Experimental flowchart of lIver fibrosis mouse model and AAV intraperitoneal injection. (B) In vivo imaging at the mouse cellular level of EGFP after AAV treatment. (C) immunofluorescence staining for SIRT4 and Desmin was performed. (D) immunofluorescence for α-SMA and GrazmB(Left panel), α-SMA and GrazmB(Right panel). (E) H&E, Masson, Sirius Red staining after AAV8-pGFAP-Sirt4 and Antii-Asialo-GM1 treatment. (F) Detecting the level of α-SMA and SIRT4 by using Western blotting analysis.

## Discussion

In the present study, we investigated the potentially protective effect of SIRT4 on liver fibrosis. SIRT4 was decreased during the activation of HSCs. To clarify the impact of SIRT4 on LF, we employed both loss-of-function and gain-of-function models, revealing that SIRT4 played a crucial role in the NK cell-mediated killing of HSCs. Mechanistically, SIRT4 promoted the expression of ULBP1 and ULBP2 via the AMPKα-pP53-NKG2DL signaling axis. In vivo, our results indicated that the deletion of SIRT4 exacerbated liver fibrosis induced by CCl4 and DDC. Furthermore, the activation of SIRT4 through targeted delivery to HSCs using AAV8 conferred protection to mice against CCl4-induced LF. Collectively, our study demonstrates that SIRT4 mitigates LF by enhancing the cytotoxic elimination of HSCs by NK cells.

NK cells play a pivotal role in the immune response within the liver, particularly through their interaction with HSCs via NKG2DLs[21]. Activated HSCs express various NKG2DLs, which serve as markers of cellular stress and activation. The recognition of these ligands by NK cells activates their cytotoxic functions, leading to the direct induction of apoptosis in HSCs[22]. This process primarily involves the release of cytotoxic granules such as perforin and granzymes, which are essential for the killing of aberrant HSCs and contribute to the resolution of liver fibrosis.Simultaneously, the engagement of NKG2DLs on HSCs triggers NK cell activation, resulting in the secretion of pro-inflammatory cytokines like IFN-γ[23]. This cytokine not only enhances the cytotoxic potential of NK cells but also modulates HSC behavior. Through IFN-γ signaling, NK cells can influence HSC activation status, thereby promoting a shift in the dynamic balance between fibrogenesis (the formation of fibrous tissue) and fibrosis resolution. The delicate interplay between NK cells and HSCs is crucial for maintaining liver homeostasis, preventing excessive fibrosis, and promoting tissue repair [24–25]. Given the significant role of this interaction in liver pathology, targeting NK cell activity and their responses may hold therapeutic promise for conditions such as liver fibrosis and cirrhosis [26]. In this study, we showed that SIRT4 enhanced the cytotoxicity of NK cells to HSCs by increasing the expression of ULBP1/ULBP2 in HSCs in vitro and in vivo.

While interrogating how SIRT4 modulates ULBP1/ULBP2 expression in HSCs, we for the first time found that SIRT4 activated the phosphorylation of p53 but had no effect on the total p53 in HSCs. Previous studies have reported that p53 can act as a transcription factor that binds to the promoter region of the genes of ULBP1/ULBP2, increasing their transcription. Meanwhile, we demonstrated that the phosphorylation of p53 could bind to the promoter region of ULBP1/ULBP2 in LX-2 cells by the method of CUT&RUN, indicating that p53 played a connecting role for SIRT4 in regulating ULBP1/ULBP2.

AMPKα plays a crucial role in the regulation of bioenergy metabolism. The activity of AMPKα is influenced by numerous factors, including glutamine metabolism[27]. Our previous studies have revealed that the inhibitory effect of SIRT4 on glutamine metabolism contributes to the activation of AMPKα. Consistently, we also confirmed that SIRT4 can inhibit glutamine metabolism, leading to the phosphorylation of AMPKα in HSCs[28]. Furthermore, in response to metabolic stress, AMPKα has been reported to induce p53 by phosphorylating and activating MDMX^28^[28]. In this study, we established a connection between SIRT4 and phosphorylated p53 via AMPKα. We demonstrated that the activation of AMPKα following SIRT4 knockdown in LX-2 cells can rescue SIRT4-induced p53 phosphorylation.

AAV is a replication-defective microvirus found in a variety of vertebrates [15]. AAV possesses a favorable safety profile and has not been associated with any human disease, establishing it as a major platform for in vivo gene therapy delivery [16]. Numerous serotypes of AAV exist, and the variability in the amino acid structures and sequences of their coat proteins, along with their interactions with host cells, has led to significant differences in infection efficiency among various AAV serotypes across different cells and tissues [17]. Literature reports indicate that AAV8 exhibits a strong targeting ability towards the liver. We constructed a liver-specific gene delivery system using AAV8 containing the promoter of glial fibrillary acidic protein (pGFAP) for targeted SIRT4 overexpression in hepatic stellate cells (HSCs) in mice. The results demonstrated that our construct, AAV8-pGFAP-SIRT4, effectively targeted HSCs. Furthermore, we confirmed the activation of NK cells in the mouse liver and the reversal of LF following viral gene injection therapy.

In summary, we have established SIRT4 as an anti-fibrogenic factor in the cytotoxic effects of NK cells on HSCs and LF. Specific targeting activates SIRT4 expression in HSCs may represent approaches to reverse LF.

## Material and Method

### Human liver samples

Samples of human liver were collected from patients at the Nanjing Drum Tower Hospital, the Affiliated Hospital of Nanjing University Medical School, who were diagnosed with fibrosis or other liver diseases, including peripheral tissues in hepatic hemangioma. This study was approved by the clinical research ethics committee of Nanjing Drum Hospital and conducted in accordance with the principles outlined in the Declaration of Helsinki. Prior to liver surgery or biopsy, informed consent was obtained for tissue analysis.

### Animal experiments

To induce liver fibrosis mouse model, C57BL/6J mice were injected with Carbon tetrachloride (CCl4) (0.75 mL/kg, Sigma-Aldrich, St. Louis, MO) twice a week for 8 weeks to induce liver fibrosis. In another model, C57BL/6J mice were fed a standard (control) diet or the standard diet supplemented with 0.1% 5-diethoxycarbonyl-1,4-dihydrocollidine (DDC; Sigma-Aldrich, Vienna, Austria) for 4 weeks.

For in vivo SIRT4 overexpression, SIRT4-His plasmids were driven by the pGFAP promoter and packaged into AAV8 for the overexpression of Sirt4 in HSCs.The mice were injected with AAV8-pGFAP-control (AAV8-pGFAP empty vector), AAV8-pGFAP-SIRT4 or (1 × 10^12^ vg/ml in a volume of 100 μl per mouse) via the tail vein two weeks after CCl4 injection. The AAV8 vectors were designed and produced by Hanbio Biotechnology Co., Ltd. (Shanghai, China). After 6 weeks, A portion of AAV8-injected mice were taken and injected intraperitoneally with Ultra-LEAF^TM^ Purified anti-Asialo-GM1 Antibody(Dakewe, Shenzhen China) in order to clear hepatic NK cells.

### Cell line treatment

The human HSC line LX-2 and the NK92 cell was purchased by purchased from Haixing Bioscience. LX-2 cells were cultured in Dulbecco’s modified Eagle’s medium (DMEM; Gibco) supplemented with 10% fetal bovine serum (FBS; Gibco).NK92 cells were cultured in a Specialised medium for NK92 cells(Haixing Bioscience). Both of them incubated at 37℃ with 5% CO2. Overexpression of SIRT4 was achieved using plasmids (#G0180974-2, Ibsbio). Transfection of SIRT4 plasmid and empty vector controls was performed using Lipofectamine 3000 (Thermo Fisher Scientific) according to the manufacturer’s instructions for 48 hours in 6-well plates. After 72 hours, the cells were treated with G418 (Selleck, MO, USA) to select for stably transfected clones.

### Lentivirus construction and infection

SIRT4 shRNAs and nonsense control shRNAs were inserted into the plasmid vector GV493, from which lentiviruses were constructed and purchased from Shanghai GeneChem Co., Ltd. Cells were infected with 8 µg/ml polybrene and subsequently selected with puromycin according to the manufacturer’s instructions. Following selection, cells were harvested for qRT-PCR, immunoblotting, and other analyses. The shRNA target sequences employed in this study were as follows: nonsense, 5′-TTCTCCGAACGTGTCACGT-3′; SIRT4, 5′-GCTGCAAGAGCGTTTCCAAGT-3′, 5′-GCGTGTCTAAACTGAATTCT-3′, 5′-GGATCATCCTTGCAGGTATAC-3′.

### Isolation and Culture of Primary HSCs

Mouse primary HSCs were isolated through collagenase digestion followed by centrifugation in an OptiPrep gradient, as previously described [29]. The cells were then cultured in DMEM supplemented with 10% fetal bovine serum (FBS) overnight, with the medium being changed every two days.

### Immunofluorescence

Cells subjected to different treatments were planted at a density of 1 × 10^3^ cells per well in 24-well plates. Seventy-two hours later, the cells were fixed for 15 minutes at room temperature using 4% paraformaldehyde and permeabilized for 15 minutes with 0.2% Triton X-100. Following a one-hour blocking period at room temperature with 5% bovine serum albumin (BSA), the cells were incubated with primary antibodies overnight at 4 °C. After three rounds of washing, the cells were cultured for one hour at room temperature in the dark with either Alexa Fluor® 488 (Abcam, ab150077) or Alexa Fluor® 647 (Abcam, 150115). Subsequently, the cells were treated for 20 minutes with DAPI (Beyotime, C1005). Finally, the cells were visualized using a fluorescence microscope (Olympus, Tokyo, Japan).

### Cytotoxicity assay

The cytotoxicity of NK cells against hepatic stellate cells (HSCs) was assessed using the lactate dehydrogenase release assay. In brief, NK cells were added to HSCs in 96-well plates at an effector-to-target ratio of 10:1, following the seeding of approximately 5 × 10^3^ HSCs per well. According to the manufacturer’s instructions, the plates were incubated for 6 hours, and the lactate dehydrogenase Cytotoxicity Assay Kit (#C0017; Beyotime) was employed to evaluate killing efficiency. The cell death ratio (%) was calculated using the formula: Cell death ratio (%) = [(Asample − Acontrol)/(Amax − Acontrol)] × 100, where A represents the absorbance value, to quantify cytotoxicity.

### Western blot analysis

Western blotting and protein isolation were done by earlier instructions [30],. The anti-SIRT4 (1:1,000, Invitrogen, # PA5-114377), anti-α-SMA, anti-collagen I, and anti-actin rabbit monoclonal antibodies (mAbs, 1:1,000; Proteintech) were the main antibodies employed. The secondary antibodies, such as goat anti-mouse IgG or goat anti-rabbit IgG conjugated with HRP (Cell Signaling Technology, MA, USA), were diluted 1:2000.

### CUT & RUN assay

CUT & RUN assay was conducted following the manufacturer’s protocol (Vazyme, HD101). Briefly, PBS or 500 mM IPA-treated Lx2 cells were incubated with ConA Beads Pro at room temperature for 10 min, anti-p53 antibody (1:50, CST, A22264) was added and rotated at room temperature for 2 h, then washed twice, added pG-MNase Enzyme and incubated at 4 C for 1 h, then washed twice, Cacl2 was added and incubated for 1 h on ice, added stop buffer and incubated at 37 C for 30 min. DNA was extracted and qPCR was used to detect pP53 binding to the DNA promoters of ULBP1 and ULBP2. A spike in DNA derived from the DNA of coli was used for uniform correction. The primer sequences used are shown in the Supplementary Material: Supplementary Table S1.

### Luciferase assay

293T cells were cotransfected with ULBP1 or ULBP2 firefly luciferase reporter plasmid (GenePharma, Shanghai, China) and TP53-WT, or an empty vector plasmid (500ng/well) using Lipofectamine 3000. Reporter assays were performed 72 h after transfection. Luciferase activity was tested with a Double-Luciferase Reporter Assay Kit(TransGen Biotech, Beijing, China) using the Dual-Light Chemiluminescent Reporter. Gene Assay System (Berthold, Germany) and was normalized to Renilla luciferase activity.

### Quantitative real-time PCR

Total RNA was extracted from the livers and HSCs using TRIzol reagent (TaKaRa, Kusatsu, Japan) according to the manufacturer’s specifications. Reverse transcription-PCR and real-time quantitative PCR analyses were performed as described by Zhang et al [31]. The primer sequences used are shown in the Supplementary Material: Supplementary Table S2.

## STATISTICS

All quantitative data are expressed as mean ± SEM. All studies were repeated at least three times with similar results. Student t-test was used to compare the two groups. One-way ANOVA followed by the Dunnett post-hoc test was used for multiple comparisons (GraphPad Software). P < 0.05 was considered statistically significant.

## Contributors

Huan Chen: Writing – original draft, Visualization, Software, Resources, Methodology, Formal analysis, Data curation, Conceptualization.

Binlin Da: Methodology, Data curation.

Zihao Cai: Software, Formal analysis, Data curation. Lei Wang: Data curation, Resources

Rui Fang:Data curation, Resources

Xiaolin Xie:Data curation, Resources

Han Zhang:Data curation, Resources

Sizhao:Data curation, Resources

Ming Zhang:Data curation, Resources

Bing Xu: Data curation, Resources

Yuzheng Zhuge: Writing – review & editing, Writing – original draft.

Feng Zhang: Writing – review & editing, Writing – original draft, Project administration,

Funding acquisition. Each author read, approved, and made changes to the manuscript’s final draft.

## Acknowledgements

This work was supported by National Natural Science Foundation of China (No. 82370628) and Jiangsu Provincial Medical Innovation Center(CXZX202213).

## Data Sharing Statement

Comprehensive data are contained within the Article. Requests for additional information should be directed to Feng Zhang and Yuzheng Zhuge

---

**Supplementary Table 1.**
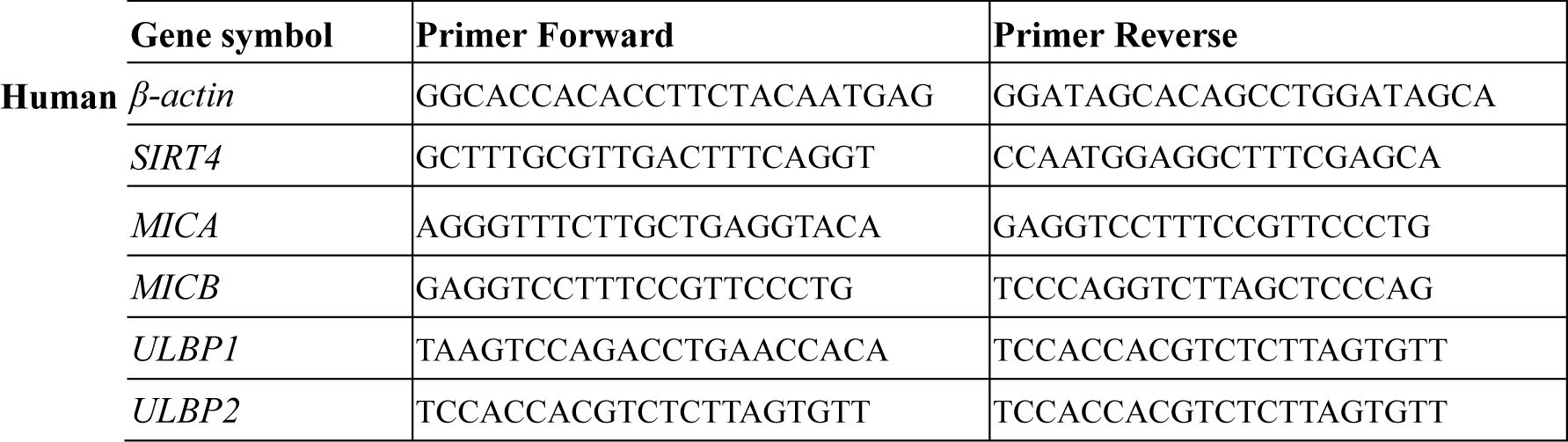
Primer used for qRT-PCR Human.

**Supplementary Table 2.**
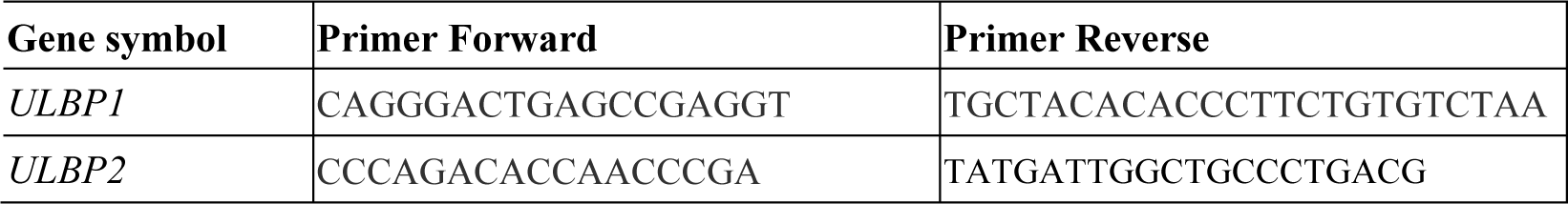
Primer used for Cut-tag-QPCR.

